# Individual Ants Do Not Show Active-rest Rhythms in Natural Nest Conditions

**DOI:** 10.1101/2020.09.17.302752

**Authors:** Haruna Fujioka, Masato S. Abe, Yasukazu Okada

## Abstract

Circadian rhythms, which respond to the day/night cycle on the earth, arise from the endogenous timekeeping system within organisms, called the biological clock. For accurate circadian rhythms, endogenous oscillations are synchronized to light and temperature. In social insects, both abiotic and biotic factors (i.e., social interactions) play a significant role in active/rest rhythm regulation. However, it is challenging to monitor individual active-rest rhythms in a colony because of the large group size and small body size. Therefore, it is unclear how social interactions coordinate each individual’s active/rest rhythms. This study developed an image-based tracking system using 2D barcodes for <I>Diacamma</I> sp. (a monomorphic ant) and measured the locomotor activities of all colony members under laboratory colony conditions. We also investigated the effect of broods on active/rest rhythms by removing all broods under colony conditions. Active/rest rhythms appeared only in solitary ants, not under colony conditions. In addition, arrhythmic active-rest rhythms were retained after brood removal. Therefore, a mixture of social interactions, not abiotic factors, induces the loss of active/rest rhythms. These results contribute to the knowledge of the diversity pattern of circadian rhythms in social insects.

## Introduction

Most organisms show daily changes in various behaviors (Saunders, 2002), such as locomotor activity (Cymborowski, 1973), feeding activity (Frisch & Aschoff, 1987), mating (Groot, 2014), and egg hatching (Itoh & Sumi, 2000). In addition, physiological phenomena, such as body temperature (Refinetti et al., 1992), blood pressure (Floras et al., 1978), endocrine hormone release (Ungar & Halberg, 1962), and metabolic activity (Chew et al., 1965), fluctuate throughout the day (Glass, 2001). This daily rhythm is not a simple response to alternating daily changes in the external environment; instead, it has an endogenous basis. When organisms are settled in constant photoperiodic and temperature conditions (i.e., “free-run” conditions), circadian rhythms (~24 h cycles) arise from the endogenous timekeeping system of the organism, called the “biological clock.” Circadian rhythms allow the organism to anticipate and prepare for changes in physical states, depending on the day-night cycle (Sharma, 2003). For accurate 24 h circadian rhythms, endogenous oscillations are synchronized to external cues (*Zeitgebers*, or time givers), such as light, temperature, and feeding time (Aschoff, 1960; Refinetti & Piccione, 2005). However, little is known about how biological factors, such as ecological interactions within and among species, affect behavioral and/or circadian rhythms.

Studies have reported on social synchronization by which animals adjust their circadian rhythms in response to social signals (Eban-Rothschild and Bloch, 2012a; Favreau et al., 2009; Regal and Connolly, 1980). To increase mating opportunities and cooperative behaviors, especially in the absence of time cues, animals require synchronization of individual active–rest cycles (Davidson and Menaker, 2003; Eban-Rothschild and Bloch, 2012b; Regal and Connolly, 1980). Interactions among mature individuals are an important time giver for cavity-dwelling and social animals (Eban-Rothschild and Bloch, 2012b; Fuchikawa et al., 2016; Mistlberger and Skene, 2004). In addition, in humans, other mammals, and birds, parents lose their daily rhythms during the reproductive phase because of interactions with offspring (Bulla et al., 2016; Lyamin et al., 2005; Nishihara et al., 2002). Similarly, bee and ant nurses lose their daily rhythms (Bloch et al., 2013; H. Fujioka et al., 2017; Moore et al. 1998). Therefore, parent-offspring interactions are one of the strongest and most important social interactions in human and nonhuman animals for changing daily rhythms.

Social insects have “arrhythmic” subterranean habitats and highly sophisticated social life; therefore, they are an excellent model for investigating the effects of environmental and social factors on daily rhythms. In addition, individuals within a colony engage in several tasks, such as foraging and brood care, that depend on the variety of social contexts and need social interactions. The circadian rhythms of insects highly depend on social contexts. For example, foraging bees typically have strong circadian rhythms regardless of their tasks, while queens and nurse bees show around-the-clock activity for egg-laying and brood care (Moore et al., 1998; Sharma et al., 2004; Shemesh et al., 2007, 2010).

In ants, in addition to the brood effect on workers’ active-rest rhythm, the behavioral rhythm seems to be highly sensitive to worker-worker interactions, too. When individual workers of *Diacamma* sp. are isolated from a colony reared under a light-dark cycle and individual behavioral rhythms measured under constant dark conditions, workers show clear active-rest rhythms regardless of age (Fujioka et al., 2017, 2019). Experimentally manipulating worker-worker interactions leads to decreased active-rest rhythms in small groups (up to five) of old (forager-age) workers (Fujioka et al., 2019). In contrast, young (nurse-age) workers show clear active-rest rhythms when five ants are grouped together, but the grouping with brood or old workers decreases active-rest rhythms. The results indicate that individual ants harbor circadian rhythms, but the expression of behavioral rhythms varies depending on the castes and the stages of interacting opponents. So far, the variability of rhythmic states in solitary and small ant groups with experimentally controlled interactions has been determined in *Diacamma* sp. However, it is important to know about individual active-rest rhythms under natural colony conditions and how such active–rest rhythms are integrated into colony-level activities.

In social insects with a large colony size, it is not easy to monitor individual active–rest rhythms under colony conditions. Colony-level observations have focused on foraging activity, oxygen consumption, and temperature, but a few studies have addressed separate individuals’ activities (Jürgen Stelzer et al., 2010b; Moritz and Kryger, 1994). Recent developed digital tools help us to automatically monitor ant behavior in groups (Greenwald et al., 2015; Mersch et al., 2013). In this study, we developed a tracking system using 2D barcodes for *Diacamma* sp. from Japan and measured all colony members’ locomotor activities under colony conditions. We also examined the effect of social interactions on active–rest rhythms. From our observations of small group interactions (Fujioka et al., 2019), we hypothesized that individuals lose active–rest rhythms under colony conditions because of a mixture of interactions with the brood, between young and old workers, and within old workers. To test this hypothesis, we compared individual activity rhythms under colony conditions with those under solitary conditions and then analyzed the effect of broods on activity rhythms by removing all broods.

## Materials and Methods

### Insect and colony setup

The ponerine ant *Diacamma* sp. from Japan (the only species of genus *Diacamma* found in Japan) is a monomorphic ant. We excavated 16 colonies in Sueyoshi Park (Naha), the Tamagusuku Youth and Children’s Center (Tamagusuku), and Kenmin-no mori (Onna), Okinawa, Japan and maintained them in the laboratory in artificial plastic nests filled with moistened plaster. To minimize environmental effects (i.e., variability of receiving light intensity), we kept the experimental colonies under a 16/8 h day–night cycle from 7:00 a.m. to 11:00 p.m. at 25°C and then transferred them to measurement conditions with constant dim-red light (i.e., imitating the all-dark [DD] free-run condition) inside the nests by covering the top of the nests with a red film. Reared colonies were fed chopped mealworms and crickets thrice a week. Each colony comprised a single queen and workers. Workers 0–30 d old stayed inside the nest and cared for the brood (i.e., eggs, larvae, and pupae), while workers older than 30 d started to forage outside the nest for various terrestrial invertebrates (Nakata, 1995). The lifespan of individuals was ~1 year (Tsuji et al., 1996). To investigate the relationship between rhythmicity and age, we marked newly eclosed workers using enamel paint (TAMIYA, Japan) at least once a month to record each individual’s age (Figure 1a).

**Figure 1.**
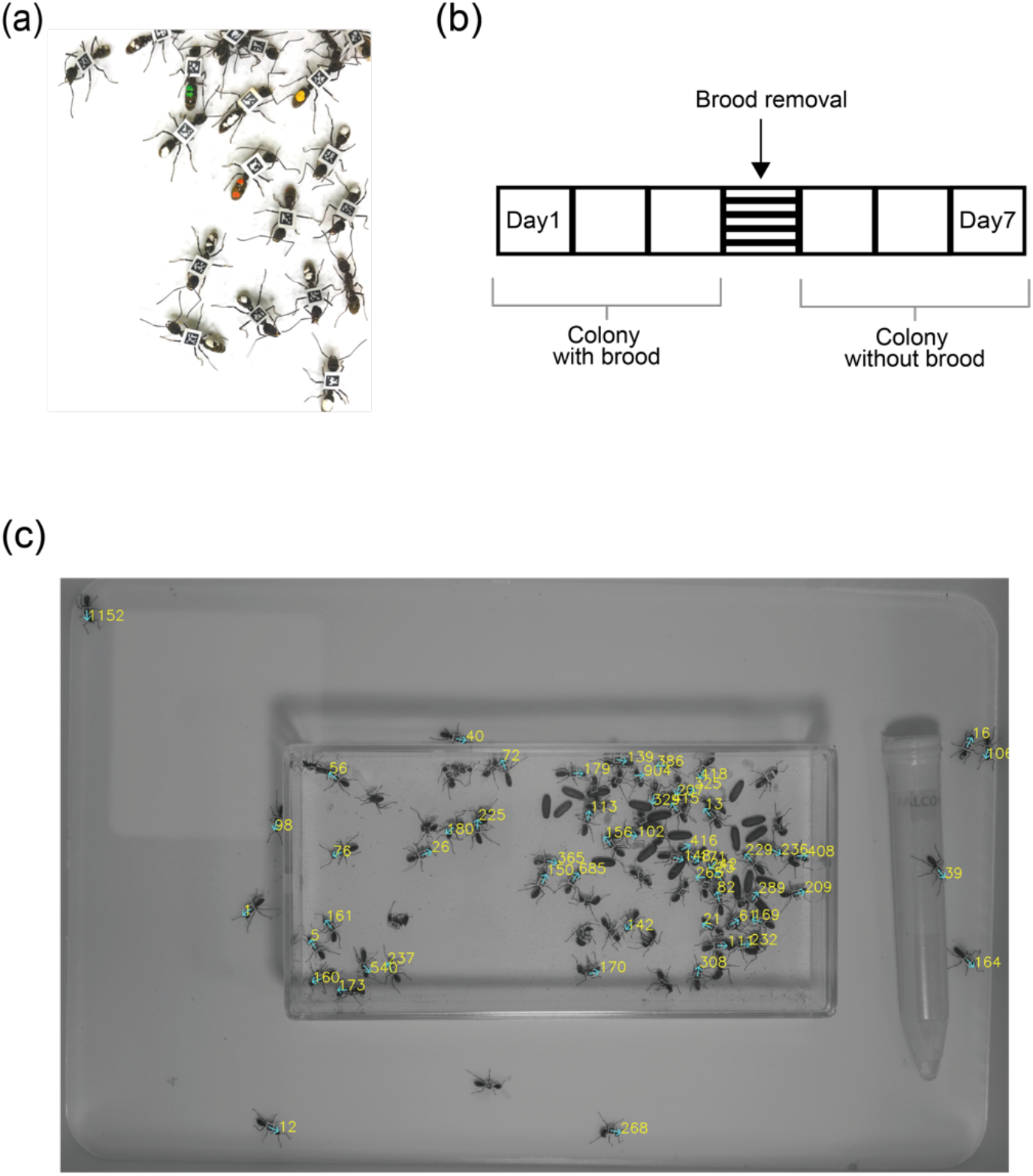
Experimental outline and 2D barcode tag–based tracking system (a) All brood (eggs, larvae, and pupae) were removed on day 4 of the experiment from two colonies. 2D barcode tags (2.0 × 2.0 mm^2^) were attached to each individual. (b) We studied two colony conditions: colony with the brood (days 1–3) and without the brood (days 5–7). (c) Experimental setup. The tag-based tracking system detected coordinates (*x, y*) and a direction of each individual every second. The square plastic box at the center is an artificial nest that was filled with moistened plaster.

### Experimental conditions

We studied three conditions: (1) solitary (without the brood), (2) colony with the brood (colony control), and (3) colony without the brood (brood removal). Workers less than 30 d old were defined as young workers (YWs) or nurse-age workers, while those older than 30 d were defined as old workers (OWs). Under solitary conditions, we randomly selected one individual from each colony and measured its locomotor activity.

Condition 1 (solitary condition) data that were different from data under conditions 2 and 3 (colony conditions) were obtained from the colonies (*n* = 8). Since the removal of a portion of workers from a colony might cause behavioral changes in the rest of the workers, we avoided this potential confound effect by using different colonies.

We used naive colonies under condition 2 (*n* = 8 colonies [A–H]), and some of them (*n* = 2 colonies) were subjected to brood removal treatment (condition 3). Under condition 2, we investigated individuals’ active–rest rhythms under laboratory conditions. All eight colonies had a queen, workers, and the brood (eggs, larvae, and pupae) each. We conducted 3 d of recording (Figure 1b). During the observation period, the ants were fed one or two crickets at the same time every day (colonies A–B: 1800; colonies C–H: 1100), and water was provided at all times. Each colony comprised 88– 194 individuals, including the queen, but we could not obtain data of some individuals because of the loss of tags, death, or tag detection insufficiency (see the “Data Processing and Checking Errors” section). Table 1 shows other detailed information about colonies.

**Table 1.**
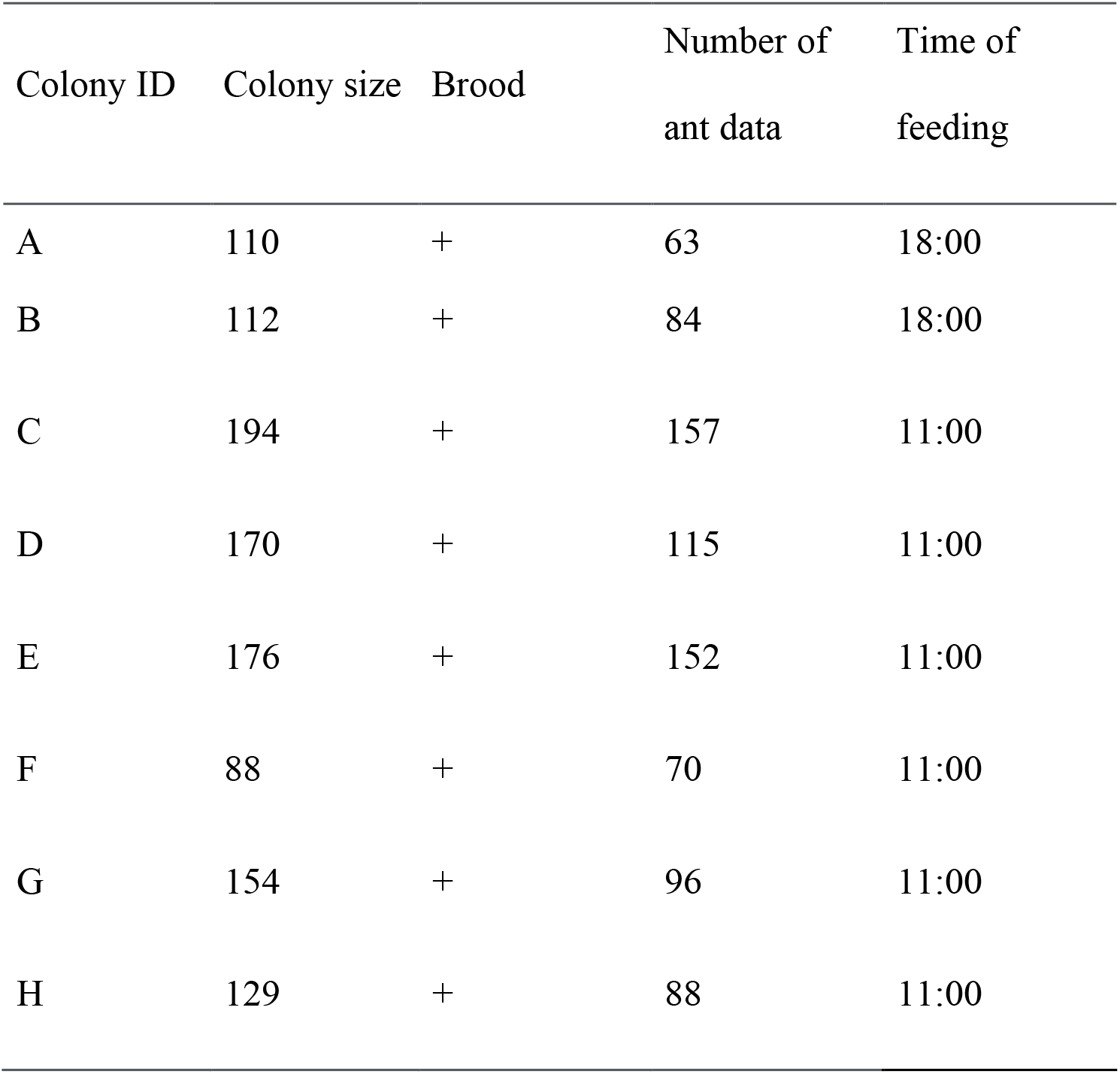
Colony information: The colony size is the number of individuals in the colony. The brood includes eggs, larvae, and pupae. We fed the ants at the same time every day.

| Colony ID | Colony size | Brood | Number of<br>ant data | Time of<br>feeding |
| --- | --- | --- | --- | --- |
| A | 110 | + | 63 | 18:00 |
| B | 112 | + | 84 | 18:00 |
| C | 194 | + | 157 | 11:00 |
| D | 170 | + | 115 | 11:00 |
| E | 176 | + | 152 | 11:00 |
| F | 88 | + | 70 | 11:00 |
| G | 154 | + | 96 | 11:00 |
| H | 129 | + | 88 | 11:00 |

Under condition 3, we used two of eight colonies under condition 2. After 3 d of recording of colonies under condition 2, we removed all broods (eggs, larvae, and pupae) on day 4 under the dim-red light from colony A and B. From day 5, we started 3 d of recording of brood removal treatment (Figure 1b). The colonies were kept under constant dim-red light throughout the experiment. To investigate the effect of broods, we compared rhythmicity and total activity between colonies under condition 2 and 3.

### Recording and analyses of activity rhythms in colonies

Condition 1 was identical to the experimental setup of Fujioka et al. (2017). Briefly, we introduced one individual into an 8 × 8 cm^2^ square plastic container and continuously recorded its locomotor activity for 3 d (60 s × 60 min × 24 h × 3 d) at 25°C ± 1°C under dim-red light (DD). We recorded the experimental area using a Logicool HD Pro Webcam C920t web camera (Hitachi, Ibaraki, Japan) and used a home-made code based on the OpenCV computer vision library (Intel Corporation, mountain View, CA, USA), to track each individual’s position. We also obtained the trajectory of movement (*x*(*t*), *y*(*t*)) every second.

Under conditions 2 and 3, we applied the image-based tracking system with tags to identify individuals. To do so, we attached a 2.0 × 2.0 mm^2^ BugTag 2D barcode tag (Robiotech, Hollywood, MD, USA) to each individual’s dorsal thorax using multipurpose, instant liquid glue (Product No. 7004; 3M, Saint Paul, MN, USA) (Figures 1a and S1); the tags and glue had no effect on mortality (Table S1; Abe S. and Fujioka 2017). We photographed the nest and foraging area every second using a Grasshopper®3 charge-coupled devide camera (GS3-U3-123S6C-C, 3000 × 4000 pixels; FLIR Systems, Wilsonville, OR, USA) for 3 d (60 s × 60 min × 24 h × 3 d) at 25°C ± 1°C under constant dim-red light. Barcode-labeled individuals were identified using a BugTag commercial vision-based tracking system (Robiotech) (Figure 1c). We again obtained the trajectory of movement (*x*(*t*), *y*(*t*)) every second (Figure 1c). Experiment nests were filled with moistened plaster (20.0 × 10.0 × 5.0 cm^3^), and each nest had one entrance of 1.0 cm diameter to allow the ants to forage (37.0 × 25.0 × 11.5 cm^3^ (Figures 1c and S2).

### Data processing and checking errors

Unfortunately, we could not track some individuals because of the detachment of tags or death; these individuals were omitted from analysis. For others, data with missing tag detection time (>50%) were also omitted from analysis. Therefore, the number of individuals analyzed was different from the actual colony size (Table 1). The reasons included image resolution (4000 × 3000) and frame rate (1 fps) for a large tracking area. In addition, focal individuals had long legs and long antennae that sometimes covered other individuals’ tags, decreasing the tag detection rate. In addition, *Diacamma* sp. often shows self-grooming behavior with a sideways posture, which prevented dorsal tag detection (H. Fujioka., personal observation).

We randomly selected 1443 discontinuous data points and manually checked the maximum movement velocity (distance/time = 36.7 mm/s). On the basis of this maximum movement velocity, we decided a threshold of 40.0 mm for excluding tracking errors. If the velocity was >40.0 mm/s, we deemed the point an error and corrected the coordinate (i.e., tag location) as not applicable (NA) data; the mean of error rates of all data was 1.39%. After processing, we filled the NA data at time *t* with previous coordinates (*t* − 1) as each individual stayed in the same position.

### Statistical analysis

All statistical analyses were performed in R v.3.5.0 and used data taken every minute. We performed *χ*^2^ periodogram analysis for the circadian active–rest rhythm (Fujioka et al., 2017; Sokolove and Bushell, 1978). The active–rest rhythm power was defined as the maximum difference between *χ*^2^ and the significance threshold line at *p* = 0.05 (Fujioka et al., 2017; Klarsfeld, Leloup, and Rouyer, 2003). The power was high when the active–rest rhythms were strong; in contrast, low or negative power indicated a weak or statistically arrhythmic state, respectively. The total activity was defined as the total movement distance during the recording period (3 d). We performed Steel–Dwass test to compare conditions 1, 2, and 3 (Figure 2). We also performed one-way analysis of variance (ANOVA) and a paired *t*-test when data followed a normal distribution with homogeneous variance (Bartlett test). In addition, we analyzed the difference of age among colonies with ANOVA. We used generalized linear mixed-effects models to test the effects of age and colony size on power or total activity. The response variables were power (a continuous value representing the active–rest rhythm strength) and total activity, while the explanatory variables were age and colony size. Linear regression models were performed to evaluate the effect of age on power and total activity.

**Figure 2.**
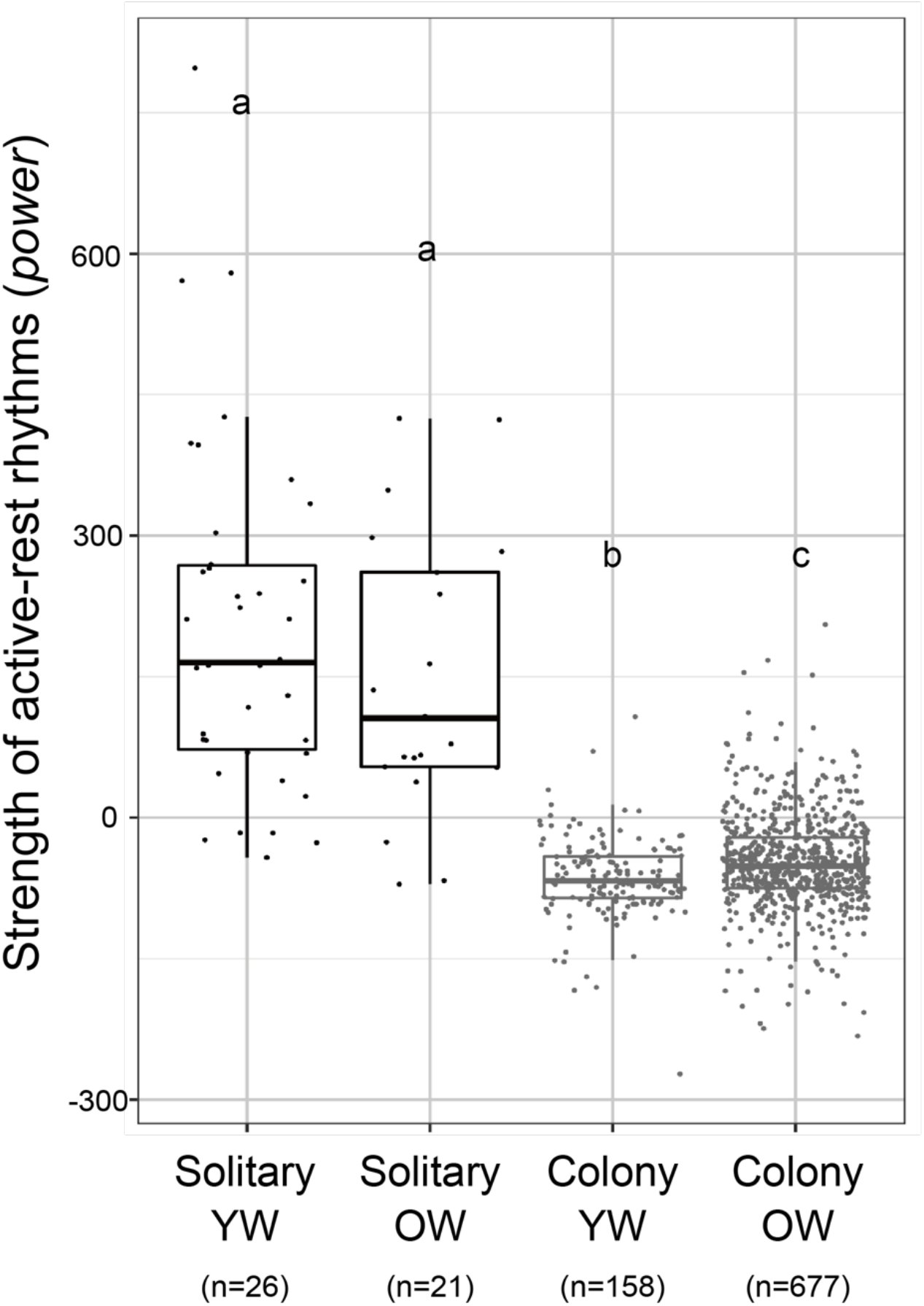
Activity rhythm under conditions 1 (solitary) and 2 (colony with the brood). Both YWs (<30 d) and OWs (>70 d) under condition 2 do not show active–rest rhythms, while YWs and OWs under condition 1 show clear active–rest rhythms. The sample size is shown below the label of the *x* axis. Different letters above the boxes indicate significant differences (Tukey–Kramer test). OW, old worker; YW, young worker.

## Result

Under condition 1, the power of most YWs and OWs was > 0, indicating that both YWs and OWs shows active–rest rhythms (Figure 2). In contrast, under condition 2, individuals almost completely lost active–rest rhythms (Figure 2; *p* < 0.01). Arrhythmic patterns were observed in the time series of both individual- and colony-level activities (Figure 3). Although there was no significant difference between YW and OW powers under condition 1 (*p* = 0.96), the OW power was greater than the YW power under condition 2 (*p* < 0.01).

**Figure 3.**
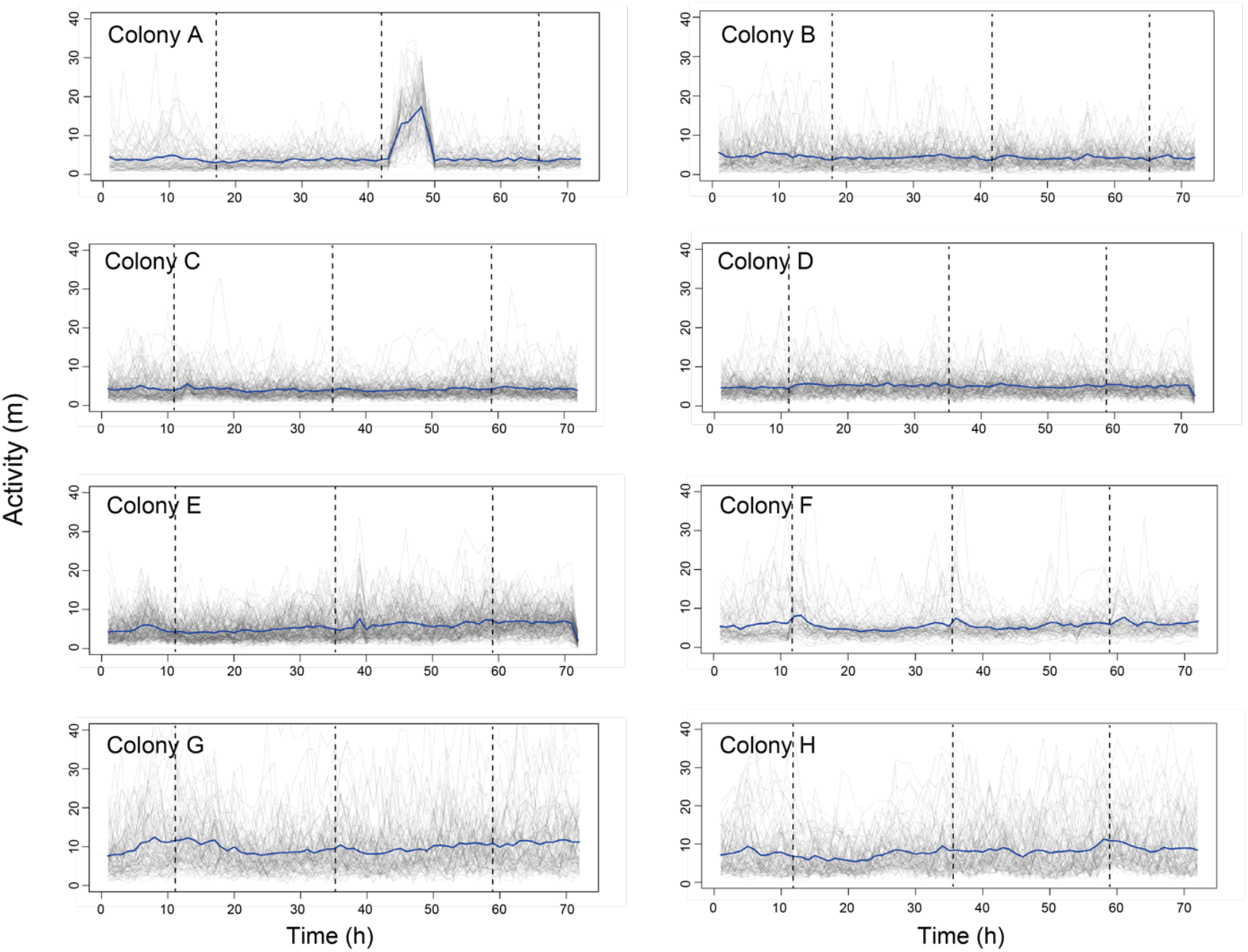
Time-series for colony- and individual-level activities for 3 d. The *x* axis represents time (h), and the *y* axis depicts the activity level (m/h). Thick (blue) and thin (gray) lines represent colony- (i.e., activity averaged over all individuals) and individual-level activity, respectively. Dashed vertical lines indicate feeding times (1800: colonies A–B; 1100: colonies C–H)

Under condition 2, power was significantly affected by age but not by colony size (Figure S3a and Table S2, age: *p* < 0.01; colony size: *p* = 0.64), while activity was strongly affected by both age and colony size (Figure S3b and Table S2; age: *p* < 0.01; colony size: *p* < 0.01). Midsize colonies showed high activity (Figure S3b). The worker age significantly differed across colonies (*F*_(7, 819)_ = 27.3; *p* < 0.01), so we separately analyzed the colonies to detect the effect of age on behaviors. In three of the eight colonies, we found a significantly negative relationship between power and age (Figure 4 and Table S3) and a positive relationship between total activity and age in seven colonies (A–C, E–H) except colony D (Figure 5 and Table S3).

**Figure 4.**
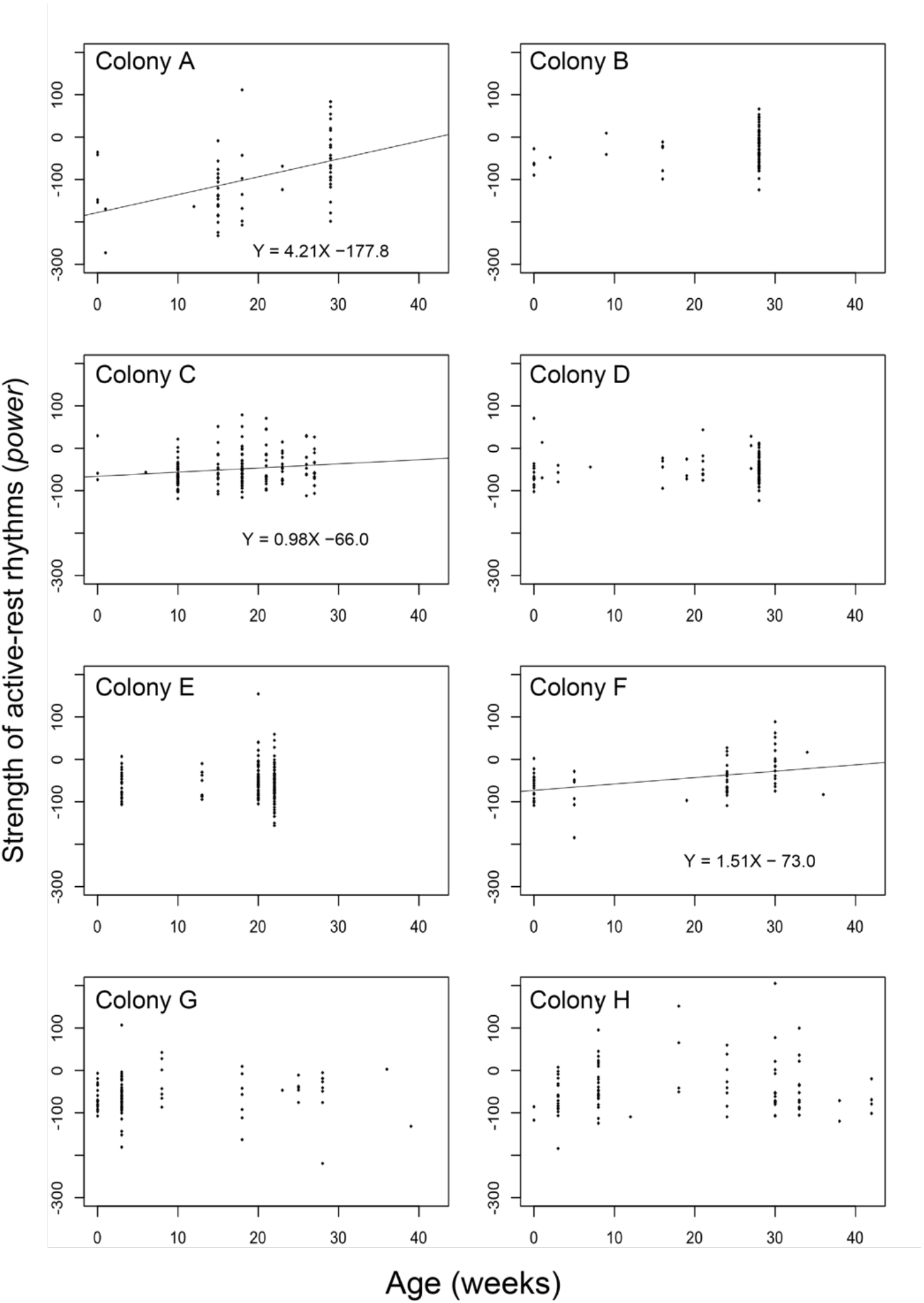
Relationship between the power (strength) of active–rest rhythms and age (weeks) in colonies A–H. The *x* and *y* axes represent the circadian rhythm power and age (weeks), respectively. The regression line represents the statistically significant relationship between power and age.

**Figure 5.**
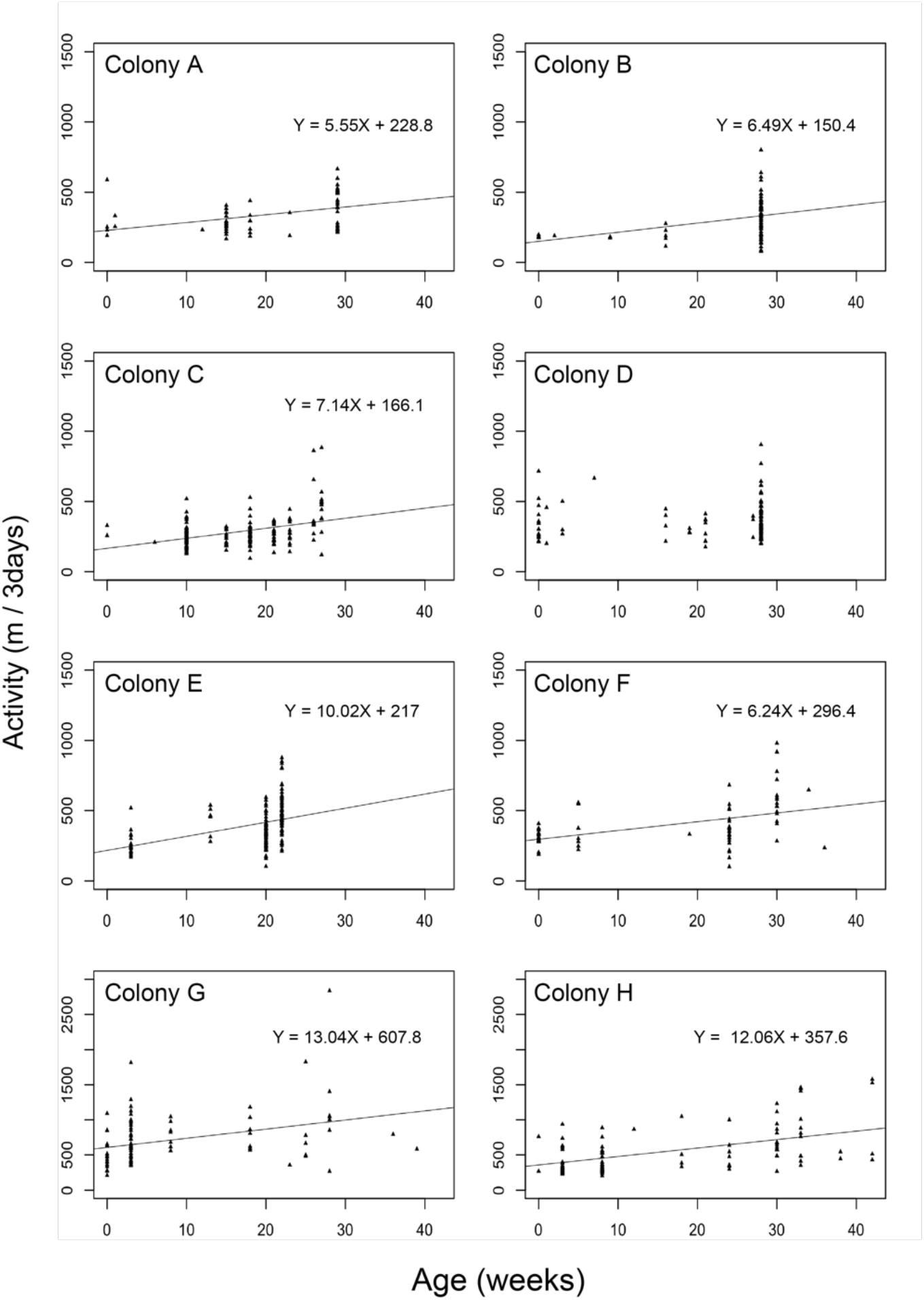
Activity levels increase with age. The *x* and *y* axes represent the total activity throughout the experimental period (m/3 days) and age (weeks), respectively. Colony D had no significant relationship between total activity and age.

Under condition 3, some individuals showed higher power, while others showed lower power after brood removal (Figure S4). On average, both power and total activity after brood removal treatment were significantly lower compared to controls (Figure 6ab). Unexpectedly, brood removal treatment still retained arrhythmic activities.

**Figure 6.**
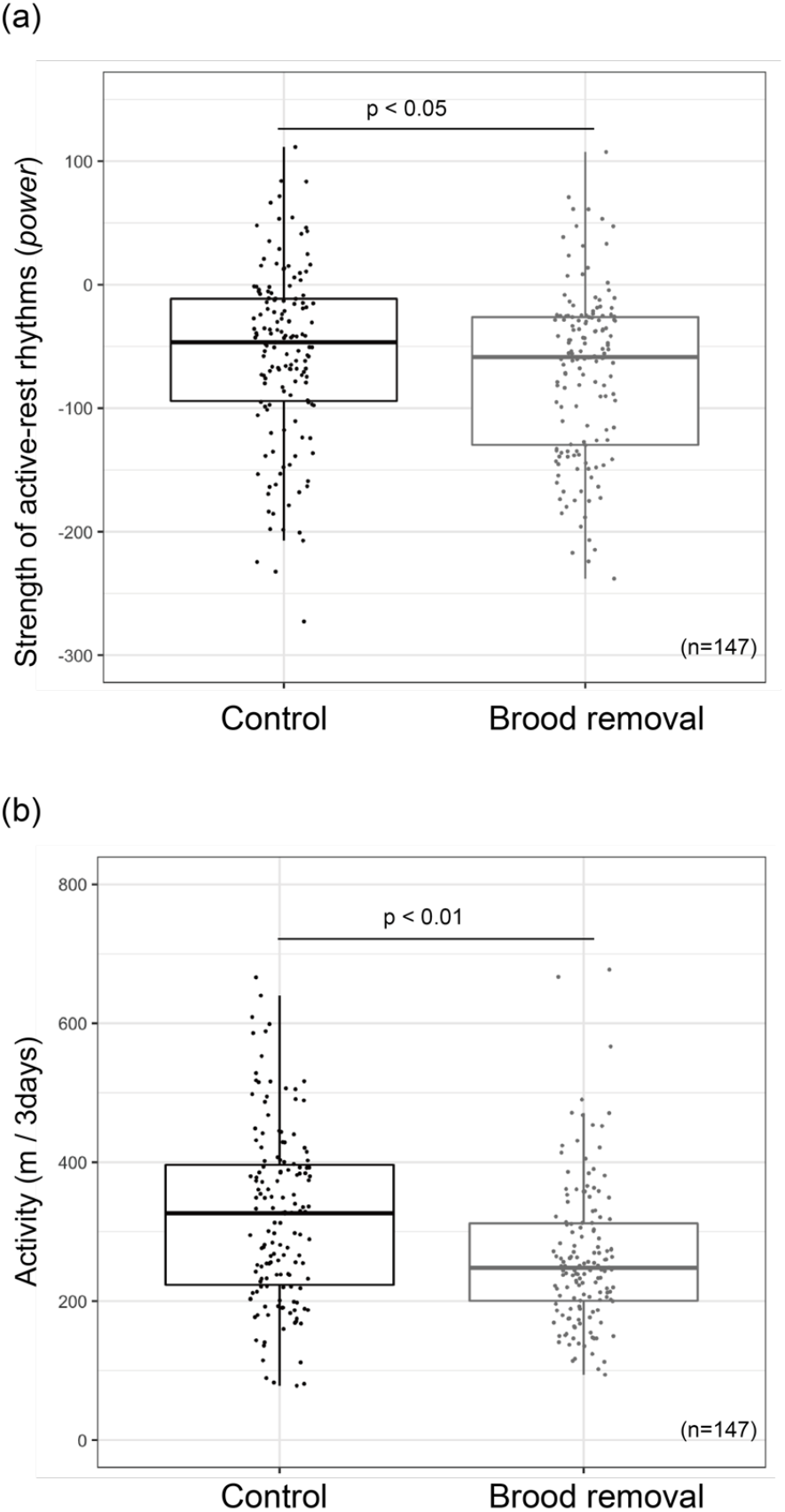
Brood removal treatment decreases power and total activity. Upper and lower panels represent the active–rest rhythm power (a) and total activity per 3 d (b). There were significant differences between control and brood removal treatment (paired *t*-test).

## Discussion

The daily rhythm is ubiquitous in organisms; however, *Diacamma* workers do not show active–rest rhythms under the seminatural colony conditions. In contrast, *Diacamma* workers show active–rest rhythms in solitary conditions, our focal individuals inherently harbor the circadian rhythm. Given the characteristic ant ecology, arrhythmic activities might be an adaptation for arrhythmic subterranean habitats and the ants’ social life. Considering the difference between the solitary and the colony conditions, we believe that social interactions, not abiotic factors, induce the loss of active–rest rhythms.

In isolated individuals, interaction with the brood decreases the rhythmicity of nursing ants (Fujioka et al. 2017). In addition, interactions with the brood or OWs decreases active–rest rhythms of YWs in small groups (Fujioka et al., 2019). Therefore, the arrhythmic activities of YWs can be attributed to interactions with the brood and/or OWs under colony conditions. Likewise, OWs show arrhythmic activities in small groups (Fujioka et al., 2019). Therefore, arrhythmic activities of OWs can be attributed to interactions with other workers under colony conditions. In laboratory-rearing nests under constant dark conditions, foragers are found all day long. Not only colony-level but also individual-level foraging activity is arrhythmic, even under colony conditions.

Our findings support the previous evidence of changing behavioral rhythms, depending on the social context (Fuchikawa et al., 2014; Fujioka et al., 2019). However, the result that foragers lose their circadian rhythms under colony conditions is inconsistent among studies. In social insects, such as bees and ants, foragers have precise daily rhythms (Bloch et al., 2001; Bochynek et al., 2017; Hoenle et al., 2019; Jürgen Stelzer et al., 2010a; Lei et al., 2019; Medeiros et al., 2014; Mildner and Roces, 2017; Moore et al., 1998; Orivel and Dejean, 2002; Passera et al., 1994; Raimundo et al., 2009; Retana et al., 1992). There are two possible reasons for this contradiction. First, previous studies did not investigate the activities of foragers inside a nest. Forager might be active during the nonforaging time inside the nest. Second, their feeding habits can affect their activities. Most bees are specialized in floral nectars and pollens, so they need to forage during flowering time. Therefore, for bees, foraging is reasonable during daytime. In contrast, temporal changes of resource availability are not straightforward for ants, which can be generalist or specialist predators, scavengers, or omnivores (feeding on nectars and preying on other arthropods) (Cerdá and Dejean, 2011; Hoenle et al., 2019). In fact, some ants, such as *Crematogaster matsumurai* (Harada, 2005), *Anoplolepis gracilipe* (Chong and Lee, 2009), *Linepithema humile* (Abril, Oliveras, & Gómez, 2007), *Diacamma* (Win et al., 2018), and *Solenopsis invicta* (Lei et al., 2019), do not have clear circadian foraging activities. Unlike the foraging time of bees, there might be no specific foraging time for generalist arthropod predators. Our focal individual feeds on various types of arthropods (Win et al., 2018). This generalist predator biology could be related to the foragers’ all-day-long activities.

Interactions with the brood (i.e. care taking) can induce arrhythmic activities in bee and ant workers (Bloch and Robinson, 2001; Fujioka et al., 2017; Nagari et al., 2019). In our focal individual, isolated and grouped nurses (up to five individuals) show circadian rhythms when freed from the brood, indicating that the presence of active–rest rhythms is the default state (Fujioka et al., 2019). Therefore, nurses might show active– rest rhythms under a brood-removed environment. The power of workers significantly decreases after brood removal. Although the increase in rhythmicity is in several YWs after brood removal treatment, this pattern is inconsistent among individuals and colonies. YWs have no or diminished rhythms because of interactions with not only the brood but also OWs (Fujioka et al., 2019). These results indicate that worker–worker interactions decrease the active–rest rhythm of YWs even when the brood is absent. One possible mechanistic explanation for the decrease in power is that brood removal treatment might change the frequency of interactions. If YW–OW interactions increase because of a loss of brood care tasks, it could eventually decrease their rhythmicity. The detailed mechanism underlying decreased active–rest rhythms requires further investigation of the interactions among workers. Besides active–rest rhythms, total activity levels also decrease after brood removal treatment. It is less likely that the decreased activity is affected by hunger. Instead, brood removal decreases the amount of work and the workers may react to the decreased workload and become inactive.

We succeeded in automatically tracking the locomotor activities of all colony members. However, it should be noted that current experimental condition (i.e. the constant temperature and dim-light) differs from the natural nest condition. Several abiotic cycles and a fluctuation of resource availability or predators exist in the wild. It is unclear how the ants share information and coordinate their active–rest rhythms within the nest. Our future research endeavor would be to understand the regulation of active–rest rhythms using manipulation experiments by controlling light and temperature.

## Conclusion

The active–rest rhythms of ants are regulated by a mixture of social worker–brood and worker–worker interactions in the colony. Multiple social interactions have non-additional, complicated effects; brood removal decreases the circadian rhythm but increases the rhythm strength. For an in-depth understanding of social chronological organization, future studies should incorporate the dynamics of social interactions. Given that ants have diverse ecological features (e.g., nesting habitats, geographical variations, diets, and foraging strategies), many more ant species might have arrhythmic colonial lives, and comparisons of different ant species should be an interesting topic. By integrating laboratory and field studies (Denlinger et al., 2017; Helm et al., 2017), the chronobiology of ants will provide new insight into the temporal organization of animals living in groups.

## Acknowledgements

Many thanks to Hakataya S. and K. Sakiyama for helping with the ant-keeping work. We also thank Uematsu J. for kindly supporting us with ant excavation.

## References

Abe S. M and Fujioka H (2017) Quantification and Analysis on Animal Social Behavior. Journal of the Robotics Society of Japan 35(6): 455–458.

Abril S, Oliveras J and Gómez C (2007) Foraging Activity and Dietary Spectrum of the Argentine Ant (Hymenoptera: Formicidae) in Invaded Natural Areas of the Northeast Iberian Peninsula. Environmental Entomology. DOI: 10.1603/0046-225x(2007)36[1166:faadso]2.0.co;2.

Aschoff J (1960) Exogenous and endogenous components in circadian rhythms. Cold Spring Harbor symposia on quantitative biology. DOI: 10.1101/sqb.1960.025.01.004.

Bloch G and Robinson GE (2001) Reversal of honeybee behavioural rhythms. Nature. DOI: 10.1038/35074183.

Bloch G, Toma DP and Robinson GE (2001) Behavioral rhythmicity, age, division of labor and period expression in the honey bee brain. Journal of Biological Rhythms. DOI: 10.1177/074873001129002123.

Bloch G, Herzog ED, Levine JD, et al. (2013) Socially synchronized circadian oscillators. Proceedings. Biological sciences / The Royal Society 280(1765). The Royal Society: 20130035. DOI: 10.1098/rspb.2013.0035.

Bochynek T, Meyer B and Burd M (2017) Energetics of trail clearing in the leaf-cutter ant Atta. Behavioral Ecology and Sociobiology. DOI: 10.1007/s00265-016-2237-5.

Bulla M, Valcu M, Dokter AM, et al. (2016) Unexpected diversity in socially synchronized rhythms of shorebirds. Nature. DOI: 10.1038/nature20563.

Cerdá X and Dejean A (2011) Predation by ants on arthropods and other animals. In: Predation in the Hymenoptera: An Evolutionary Perspective.

Chew RM, Lindberg RG and Hayden P (1965) Circadian Rhythm of Metabolic Rate in Pocket Mice. Journal of Mammalogy. DOI: 10.2307/1377637.

Chong KF and Lee CY (2009) Influences of temperature, relative humidity and light intensity on the foraging activity of field populations of the longlegged ant, Anoplolepis gracilipes (hymenoptera: Formicidae). Sociobiology.

Cymborowski B (1973) Control of the circadian rhythm of locomotor activity in the house cricket. Journal of Insect Physiology. DOI: 10.1016/0022-1910(73)90173-X.

Davidson AJ and Menaker M (2003) Birds of a feather clock together - Sometimes: Social synchronization of circadian rhythms. Current Opinion in Neurobiology. DOI: 10.1016/j.conb.2003.10.011.

Denlinger DL, Hahn DA, Merlin C, et al. (2017) Keeping time without a spine: What can the insect clock teach us about seasonal adaptation? Philosophical Transactions of the Royal Society B: Biological Sciences. DOI: 10.1098/rstb.2016.0257.

Eban-Rothschild A and Bloch G (2012a) Circadian Rhythms and Sleep in Honey Bees. In: Honeybee Neurobiology and Behavior. DOI: 10.1007/978-94-007-2099-2_3.

Eban-Rothschild A and Bloch G (2012b) Social Influences on Circadian Rhythms and Sleep in Insects. In: Advances in Genetics. DOI: 10.1016/B978-0-12-387687-4.00001-5.

Favreau A, Richard-Yris MA, Bertin A, et al. (2009) Social influences on circadian behavioural rhythms in vertebrates. Animal Behaviour. DOI: 10.1016/j.anbehav.2009.01.004.

Floras JS, Jones J V., Johnston JA, et al. (1978) Arousal and the circadian rhythm of blood pressure. Clinical science and molecular medicine. Supplement.

Frisch B and Aschoff J (1987) Circadian rhythms in honeybees: entrainment by feeding cycles. Physiological Entomology. DOI: 10.1111/j.1365-3032.1987.tb00722.x.

Fuchikawa T, Okada Y, Miyatake T, et al. (2014) Social dominance modifies behavioral rhythm in a queenless ant. Behavioral Ecology and Sociobiology 68(11): 1843–1850. DOI: 10.1007/s00265-014-1793-9.

Fuchikawa T, Eban-Rothschild A, Nagari M, et al. (2016) Potent social synchronization can override photic entrainment of circadian rhythms. Nature Communications. DOI: 10.1038/ncomms11662.

Fujioka H, Abe MS, Fuchikawa T, et al. (2017) Ant circadian activity associated with brood care type. Biology Letters 13(2). DOI: 10.1098/rsbl.2016.0743.

Fujioka H, Abe MS and Okada Y (2019) Ant activity-rest rhythms vary with age and interaction frequencies of workers. Behavioral Ecology and Sociobiology. DOI: 10.1007/s00265-019-2641-8.

Glass L (2001) Synchronization and rhythmic processes in physiology. Nature. DOI: 10.1038/35065745.

Greenwald E, Segre E and Feinerman O (2015) Ant trophallactic networks: simultaneous measurement of interaction patterns and food dissemination. Scientific reports 5. Nature Publishing Group: 12496. DOI: 10.1038/srep12496.

Groot AT (2014) Circadian rhythms of sexual activities in moths: A review. Frontiers in Ecology and Evolution. DOI: 10.3389/fevo.2014.00043.

Harada Y (2005) Diel and seasonal patterns of foraging activity in the arboreal ant Crematogaster matsumurai Forel. Entomological Science. DOI: 10.1111/j.1479-8298.2005.00115.x.

Helm B, Visser ME, Schwartz W, et al. (2017) Two sides of a coin: Ecological and chronobiological perspectives of timing in the wild. Philosophical Transactions of the Royal Society B: Biological Sciences. DOI: 10.1098/rstb.2016.0246.

Hoenle PO, Blüthgen N, Brückner A, et al. (2019) Species-level predation network uncovers high prey specificity in a Neotropical army ant community. Molecular Ecology. DOI: 10.1111/mec.15078.

Itoh MT and Sumi Y (2000) Circadian clock controlling egg hatching in the cricket (Gryllus bimaculatus). Journal of Biological Rhythms. DOI: 10.1177/074873040001500305.

Jürgen Stelzer R, Stanewsky R and Chittka L (2010a) Circadian foraging rhythms of bumblebees monitored by radio-frequency identification. Journal of biological rhythms 25: 257–267. DOI: 10.1177/0748730410371750.

Jürgen Stelzer R, Stanewsky R and Chittka L (2010b) Circadian foraging rhythms of bumblebees monitored by radio-frequency identification. Journal of Biological Rhythms. DOI: 10.1177/0748730410371750.

Klarsfeld A, Leloup JC and Rouyer F (2003) Circadian rhythms of locomotor activity in Drosophila. Behavioural Processes 64(April): 161–175. DOI: 10.1016/S0376-6357(03)00133-5.

Lei Y, Zhou Y, Lü L, et al. (2019) Rhythms in Foraging Behavior and Expression Patterns of the Foraging Gene in Solenopsis invicta (Hymenoptera: Formicidae) in relation to Photoperiod. Journal of Economic Entomology 112(6). Oxford Academic: 2923–2930. DOI: 10.1093/jee/toz175.

Lyamin O, Pryaslova J, Lance V, et al. (2005) Animal behaviour: Continuous activity in cetaceans after birth. Nature. DOI: 10.1038/4351177a.

Medeiros J, Azevedo DLO, Santana MAD, et al. (2014) Foraging activity rhythms of Dinoponera quadriceps (Hymenoptera: Formicidae) in its natural environment. Journal of Insect Science. DOI: 10.1093/jisesa/ieu082.

Mersch DP, Crespi A and Keller L (2013) Tracking Individuals Shows Spatial Fidelity Is a Key Regulator of Ant Social Organization. Science 340(6136).

Mildner S and Roces F (2017) Plasticity of daily behavioral rhythms in foragers and nurses of the ant camponotus rufipes: Influence of social context and feeding times. PLoS ONE 12(1). DOI: 10.1371/journal.pone.0169244.

Mistlberger RE and Skene DJ (2004) Social influences on mammalian circadian rhythms: Animal and human studies. Biological Reviews of the Cambridge Philosophical Society. DOI: 10.1017/S1464793103006353.

Moore D, Angel JE, Cheeseman IM, et al. (1998) Timekeeping in the honey bee colony: integration of circadian rhythms and division of labor. Behavioral Ecology and Sociobiology. DOI: 10.1007/s002650050476.

Moritz RFA and Kryger P (1994) Self-organization of circadian rhythms in groups of honeybees (Apis mellifera L.). Behavioral Ecology and Sociobiology. DOI: 10.1007/BF00167746.

Nagari M, Gera A, Jonsson S, et al. (2019) Bumble Bee Workers Give Up Sleep to Care for Offspring that Are Not Their Own. Current Biology. DOI: 10.1016/j.cub.2019.07.091.

Nakata K (1995) Age polyethism, idiosyncrasy and behavioural flexibility in the queenless ponerine ant,Diacamma sp. Journal of Ethology 13(1). Springer-Verlag: 113–123. DOI: 10.1007/BF02352570.

Nishihara K, Horiuchi S, Eto H, et al. (2002) The development of infants’ circadian rest–activity rhythm and mothers’ rhythm. Physiology & Behavior 77(1): 91–98. DOI: 10.1016/S0031-9384(02)00846-6.

Orivel J and Dejean A (2002) Ant activity rhythms in a pioneer vegetal formation of French Guiana (Hymenoptera: Formicidae). Sociobiology.

Passera L, Lachaud JP, Gomel L, et al. (1994) Individual food source fidelity in the neotropical ponerine ant ectatomma ruidum roger (Hymenoptera formicidae). Ethology Ecology and Evolution. DOI: 10.1080/08927014.1994.9523004.

Raimundo RLG, Freitas AVL and Oliveira PS (2009) Seasonal Patterns in Activity Rhythm and Foraging Ecology in the Neotropical Forest-Dwelling Ant, *Odontomachus chelifer* (Formicidae: Ponerinae). Annals of the Entomological Society of America. DOI: 10.1603/008.102.0625.

Refinetti R and Piccione G (2005) Intra- and inter-individual variability in the circadian rhythm of body temperature of rats, squirrels, dogs, and horses. Journal of Thermal Biology. DOI: 10.1016/j.jtherbio.2004.09.003.

Refinetti R, Nelson DE and Menaker M (1992) Social stimuli fail to act as entraining agents of circadian rhythms in the golden hamster. Journal of Comparative Physiology A. DOI: 10.1007/BF00196900.

Regal PJ and Connolly MS (1980) Social Influences On Biological Rhythms. Behaviour. DOI: 10.1163/156853980X00104.

Retana J, Cerda Xim and Espadaler Xavier (1992) Coexistence of two Sympatric Ant Species, Pheidole pallidula and Tetramorium semilaeve (Hymenoptera: Formicidae). Entomologia Generalis. DOI: 10.1127/entom.gen/17/1992/29.

Saunders DS (2002) Insect Clocks, Third Edition. Elsevier Science. DOI: https://doi.org/10.1016/B978-0-444-50407-4.X5000-9.

Sharma VK (2003) A simple computer-aided device for monitoring activity of small mammals and insects. Biological Rhythm Research 34(January 2015): 3–12. DOI: 10.1076/brhm.34.1.3.14078.

Sharma VK, Lone SR, Goel A, et al. (2004) Circadian consequences of social organization in the ant species Camponotus compressus. Naturwissenschaften. Springer-Verlag. Available at: http://repository.ias.ac.in/30777/1/344.pdf (accessed 19 January 2015).

Shemesh Y, Cohen M and Bloch G (2007) Natural plasticity in circadian rhythms is mediated by reorganization in the molecular clockwork in honeybees. The FASEB journal: official publication of the Federation of American Societies for Experimental Biology 21: 2304–2311. DOI: 10.1096/fj.06-8032com.

Shemesh Y, Eban-Rothschild A, Cohen M, et al. (2010) Molecular dynamics and social regulation of context-dependent plasticity in the circadian clockwork of the honey bee. Journal of Neuroscience. DOI: 10.1523/JNEUROSCI.1490-10.2010.

Sokolove PG and Bushell WN (1978) The chi square periodogram: Its utility for analysis of circadian rhythms. Journal of Theoretical Biology 72(1): 131–160. DOI: 10.1016/0022-5193(78)90022-X.

Tsuji K, Nakata K and Heinze J (1996) Lifespan and reproduction in a Queenless Ant. Naturwissenschaften. DOI: 10.1007/BF01141985.

Ungar F and Halberg F (1962) Circadian rhythm in the in vitro response of mouse adrenal to adrenocorticotropic hormone. Science. DOI: 10.1126/science.137.3535.1058.

Win AT, Machida Y, Miyamoto Y, et al. (2018) Seasonal and temporal variations in colony-level foraging activity of a queenless ant, Diacamma sp., in Japan. Journal of Ethology. DOI: 10.1007/s10164-018-0558-8.

