## Supplementary material for "Individual Ants Do Not Show Active-rest Rhythms in Natural Nest Conditions": supplemenrary material

Table S1. The results for generalized linear models for examining the effects of age and colony size on *power* and total activity.

| | Parameter | Estimate | Std. Error | t value | p |
|---|---|---|---|---|---|
| *Power* | (Intercept) | -70.19 | 8.64 | -8.12 | < 0.01 |
| | Age | 0.12 | 0.02 | 5.10 | < 0.01 |
| | Colony size | 0.02 | 0.05 | 0.47 | 0.64 |
| Activity | (Intercept) | 475.25 | 42.78 | 11.11 | < 0.01 |
| | Age | 0.32 | 0.12 | 2.64 | < 0.01 |
| | Colony size | -0.66 | 0.25 | -2.66 | <0.01 |

Table S2. Regression equations estimated using linear model.

| | Colony | Regression equation | $R^2$ | p |
|---|---|---|---|---|
| *Power* | Total | Y = 0.85X - 66.4 | 0.0366 | **p < 0.01** |
| | A | Y = 4.21X −177.8 | 0.1924 | **p < 0.01** |
| | B | Y = 0.98X - 54.2 | 0.0284 | p = 0.067 |
| | C | Y = 0.98X −66.0 | 0.0196 | **p = 0.0443** |
| | D | Y = 0.25 X -54.3 | -0.0018 | p = 0.373 |
| | E | Y = 0.33X -59.2 | -0.0038 | p = 0.516 |
| | F | Y = 1.51X − 73.0 | 0.16 | **p < 0.01** |
| | G | Y = 0.27X -61.0 | -0.0069 | p = 0.554 |
| | H | Y = 1.49X − 52.8 | -0.0108 | p = 0.792 |
| Activity | Total | Y = 2.29X + 374.3 | 0.006 | **p < 0.01** |
| | A | Y = 5.55X + 228.8 | 0.1536 | **p < 0.01** |
| | B | Y = 6.49X + 150.4 | 0.116 | **p < 0.01** |
| | C | Y = 7.14X + 166.1 | 0.1444 | **p < 0.01** |
| | D | Y = 0.87 X + 344.6 | -0.0039 | p = 0.456 |
| | E | Y = 10.02X + 217.0 | 0.1807 | **p < 0.01** |
| | F | Y = 6.24X + 296.4 | 0.1812 | **p < 0.01** |
| | G | Y = 13.04X + 607.8 | 0.111 | **p < 0.01** |
| | H | Y = 8.17X + 228.4 | 0.1535 | **p < 0.01** |

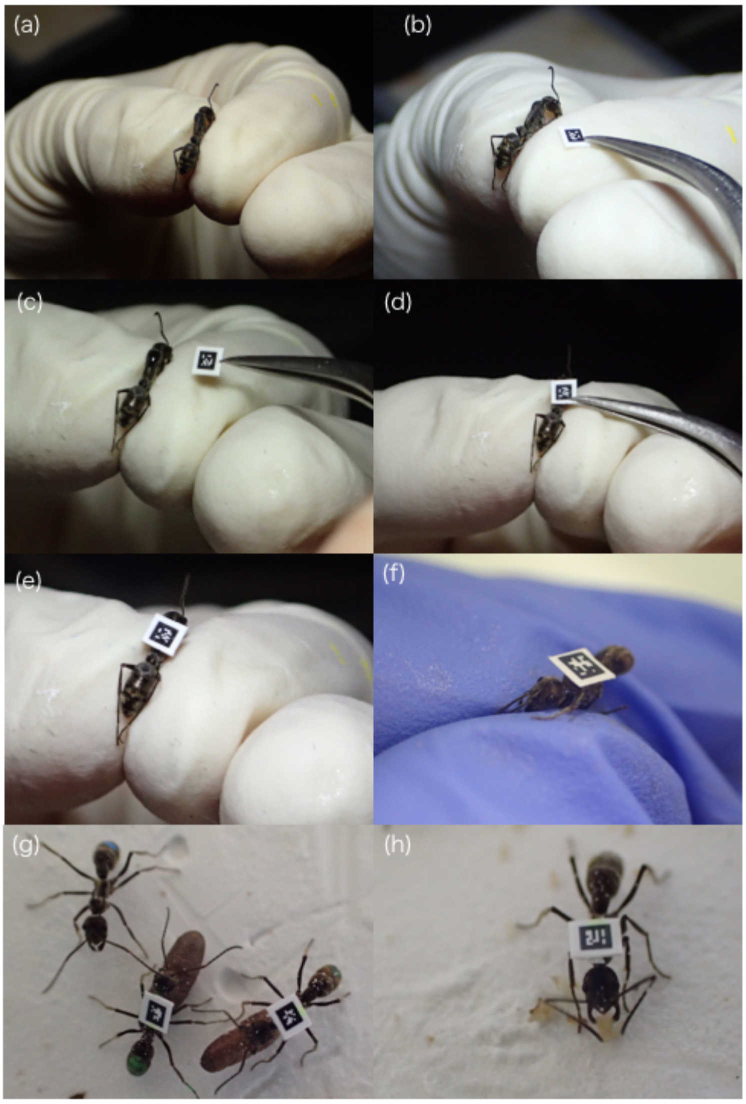


Figure S1. Tag was attached by glue without anesthesia.

We gently held ant by fingers and attached tag (a-d). We held ant for 2-3 min until the glue gets completely dry.  There is no unusual behavior in tagged ants, which performed carrying pupa (g), collecting eggs (h), grooming and dominance behaviors as well as untagged ant.

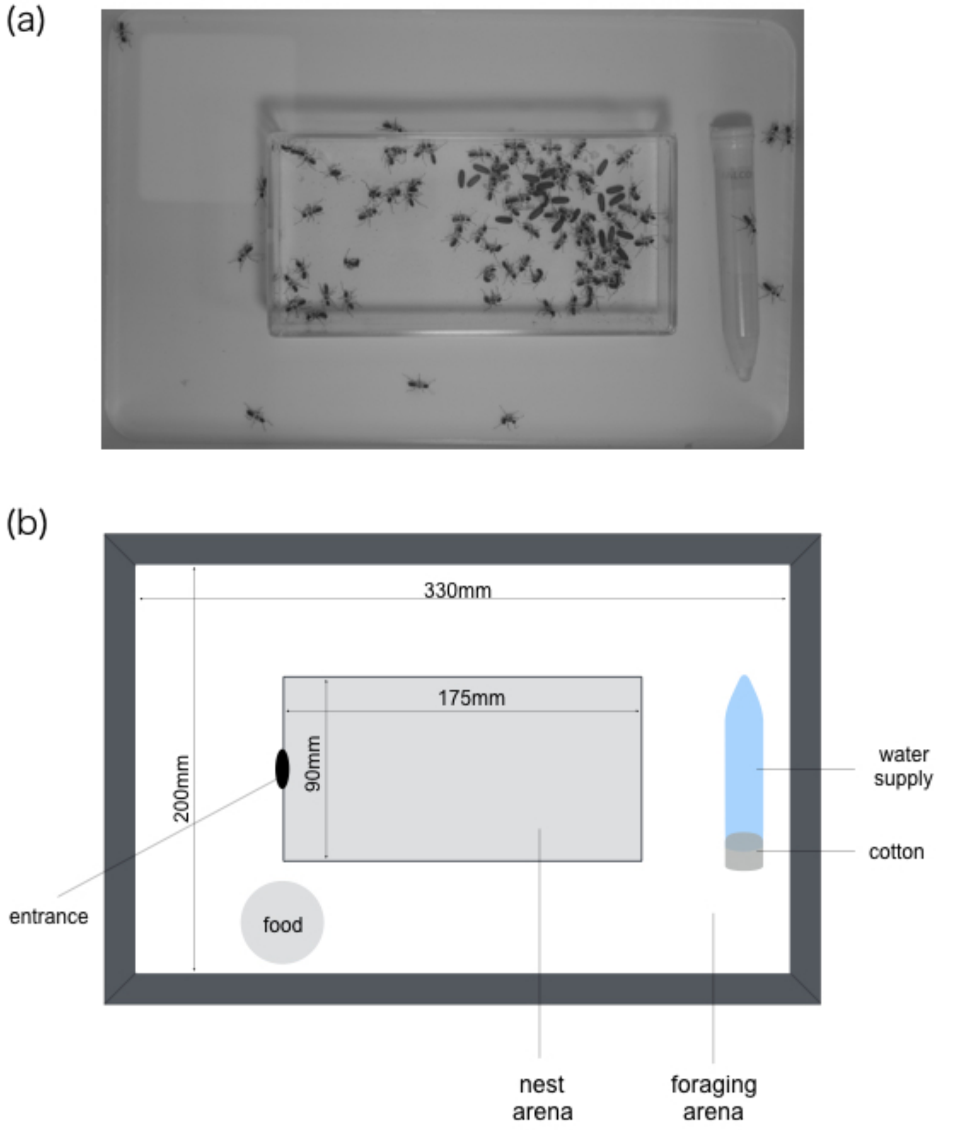


Figure S2. Colony setup during experiment. The artificial nest was a square box that were filled with a moistened plaster. Ants were prohibited from climbing the walls of nest and foraging arena by coating the lateral sides of the containers with fluoropolymer resin (PTFE-30 (Fluon), BioQuip, CA, USA).

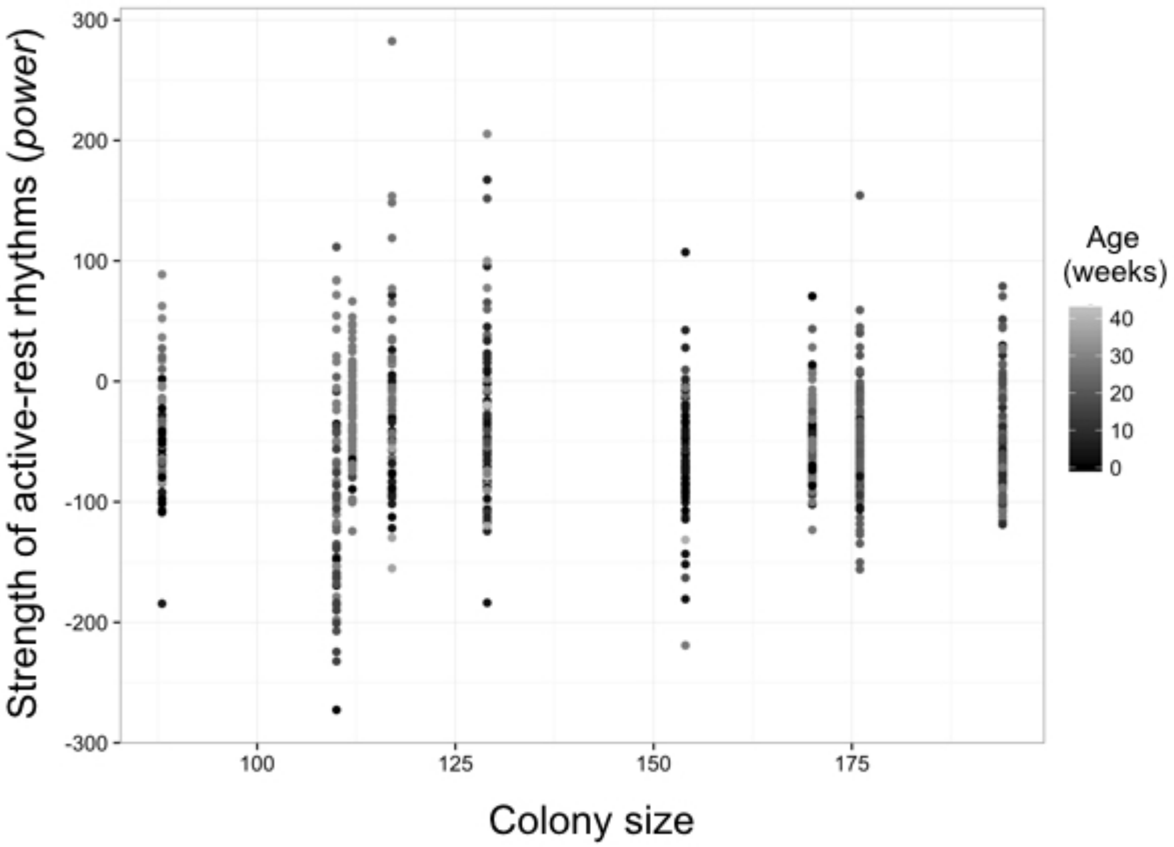


Figure S3. The effect of colony size on *power.*

Each point represents an individual. The X-axis and y-axis represent the strength of circadian rhythms (*power*) and colony size, respectively. Black is younger workers and grey is older workers.

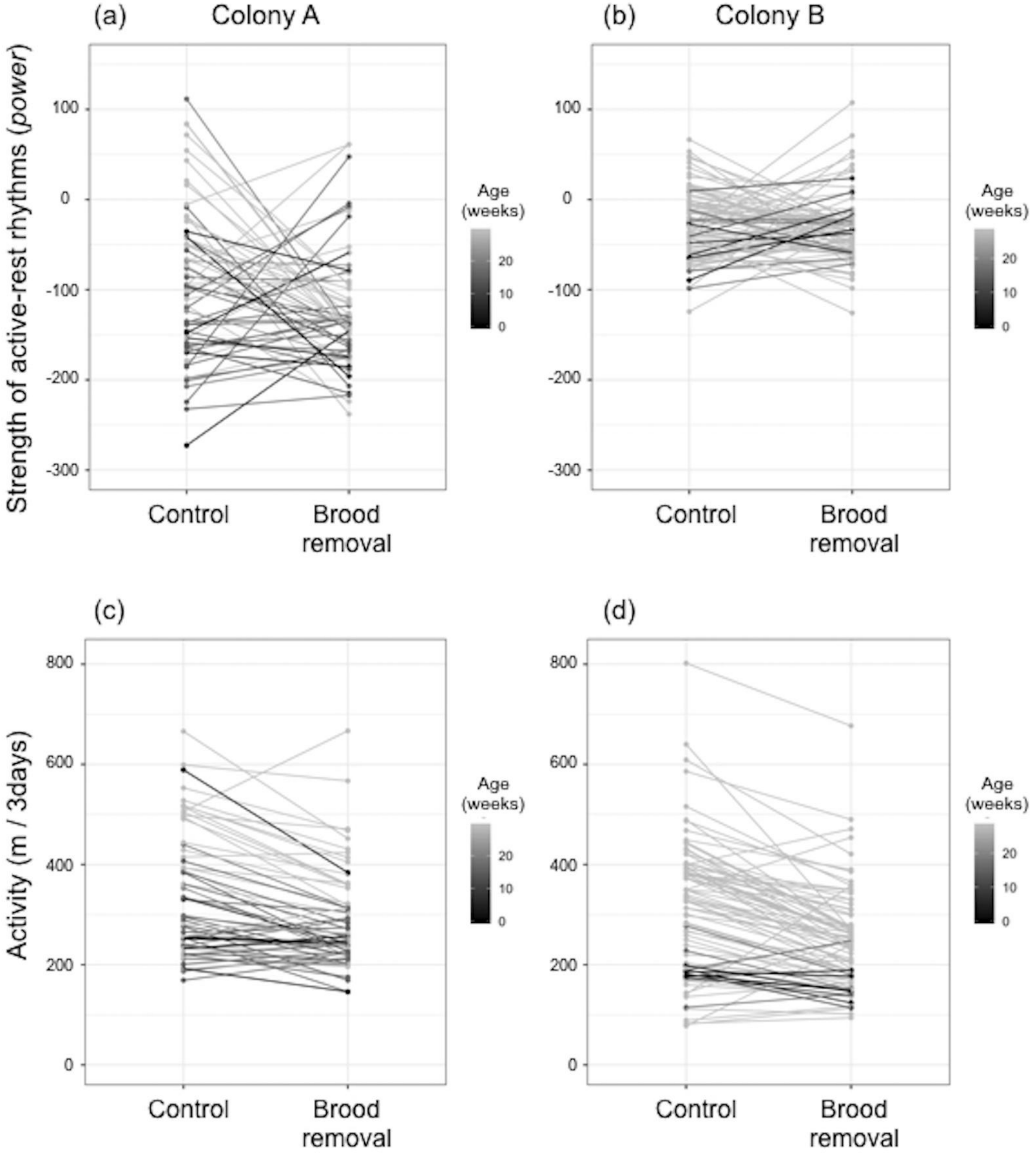


Figure S4. Brood removal decreased the level of activity.

Upper and Lower panels represent the strength of active-rest activity (*power*) (a,b) and a total of activity per 3 days (c,d). Left and right panels represent colony A and B, respectively. Color depth (black to grey) reflects age (weeks). Black is younger workers and grey is older workers.
